# Modulation of implicitly perceived hand size by visuotactile recalibration

**DOI:** 10.1101/2022.10.13.512071

**Authors:** Xaver Fuchs, Tobias Heed

## Abstract

When concurrent visual and tactile stimuli are repeatedly presented with a spatial offset, even unisensory tactile stimuli are afterwards perceived with a spatial bias towards the previously presented visual stimuli. This so-called visuotactile ventriloquism aftereffect reflects visuotactile recalibration. It is unknown whether this recalibration occurs within a bodily map and interacts with perceived features like shape and size of body parts.

Here, we applied tactile stimuli to participants’ hidden left hand and simultaneously presented visual stimuli with spatial offsets that – if integrated with the tactile stimuli – implied an enlarged hand size. We either used a fixed spatial mapping between tactile and visual positions (“congruent”), or a scrambled (“incongruent”) mapping. We assessed implicitly perceived hand size via two independent behavioral assessments: pointing movements to unisensory tactile stimuli and tactile distance judgments. Moreover, we assessed explicitly perceived change in hand size with perceptual self-reports.

Especially after congruent recalibration, participants localized unimodal tactile stimuli as if they were aiming at an enlarged hand. They also reported tactile distance as shorter after congruent than incongruent recalibration. These modulations resemble those obtained after using tools that prolong the arm and extend reaching space; they suggest that recalibration affected a common, implicit hand representation that underlies both tasks. In contrast, explicit perceptual self-reports did not differ significantly between congruent and incongruent recalibration.

Thus, simple visuotactile stimuli are sufficient to modify implicitly perceived body size, indicating a tight link of low-level multisensory processes such as the visuotactile ventriloquism aftereffect and body representation.

## Introduction

When we reach towards a touch on the skin, we must determine the target location in external space by combining skin location and body posture. Such information is often presumed to be organized in cognitive structures termed *body representations*. In addition to a conscious *body image*, Head and Holmes (1911) proposed two *body schemata*: a *superficial body schema* comprising information about where sensations are perceived on the body surface and a *postural body schema* that encompasses positional information of body parts derived from its current posture. Yet, additional information about both the size and shape of the body’s segments must be accounted for: For example, the position of the tip of the index finger, relative to the hand, can only be determined when the finger’s length is known. Accordingly, size and shape schemata have been suggested as extensions of Head and Holmes’s original proposal (Longo et al., 2010; Tamè et al., 2019).

Body perception is remarkably malleable and can be readily changed by bottom-up (multi)sensory input, sometimes within seconds (Botvinick & Cohen, 1998; Riemer et al., 2019; Ehrsson, 2007; Lenggenhager et al., 2007). In the famous rubber hand illusion (Botvinick & Cohen, 1998), participants perceive illusory touch on, and feel ownership of, a rubber hand when it is stroked in synchrony with their hidden (real) hand. Many authors have concluded that the referral of touch to a location on the rubber hand results from multisensory integration mechanisms that prioritize vision (rubber hand position) over proprioception (real hand position) and resolve visuo-proprioceptive conflict by referring the touch to the visually perceived location. An often-reported behavioral effect of the rubber hand illusion is that participants make systematic localization errors and report the perceived location of the touched hand displaced towards the rubber hand, even when vision of the rubber hand is now prevented. At the same time, they also report the rubber hand to be displaced towards the true tactile location (Erro et al., 2018; Fuchs et al., 2016). Two mechanisms have been proposed as the potential basis for such localization adjustment: Sensory recalibration pertains to the idea that the location of one sensory modality is adjusted to be similar to that of another. In contrast, reweighting pertains to the idea that the contribution of a given modality to a common, integrated location estimate is modified, resulting in a bias towards or away from that modality after assigning it more or less weight, respectively (Limanowski, 2021). These mechanisms are not mutually exclusive because they can occur simultaneously or one following upon the other (Ernst & Di Luca, 2011).

Mutual, bidirectional inter-sensory dependence of spatial perception is well-known from multisensory integration paradigms such as the ventriloquism effect (Alais & Burr, 2004) and the ventriloquism aftereffect (Recanzone, 1998), both of which refer to the modulation of perceived stimulus location in visual-auditory (Recanzone, 1998) or visuotactile cue combinations (Samad & Shams, 2018): when a tactile stimulus is presented on the forearm in synchrony with a visual flash that is spatially offset from the tactile stimulus, participants perceive the tactile stimulus displaced towards the visual stimulus (ventriloquism effect); after bimodal stimuli have been presented with an offset for some time, even *unisensory* tactile stimuli are perceptually displaced in the direction of the visual stimulus offset of the previously experienced bimodal stimulation (ventriloquism aftereffect). In line with the principle of bidirectional influence, the visual stimulus is also perceptually biased towards the auditory (Alais & Burr, 2004) or tactile stimulus (Samad & Shams, 2018), respectively. Explanations of these effects have recurred both on sensory recalibration (Ernst & Di Luca, 2011) and optimal integration (Alais & Burr, 2004), bearing obvious conceptual analogies to experiments involving bodily illusions such as the RHI (Kilteni et al., 2015).

Yet, the conceptual links between visuotactile ventriloquism aftereffects and changes of *body representation* have not been explicitly specified. For instance, it is unknown whether visuotactile recalibration affects an external spatial or rather a bodily map. Both studies on prism adaption in owls (Linkenhoker & Knudsen, 2002) and on the visual-auditory ventriloquism aftereffects in humans (Recanzone, 1998) have suggested that auditory space is recalibrated by realigning the auditory with the visual spatial map. But how would this work in touch? One possibility is that tactile stimuli are realigned with visual space and therefore perceived at a modified location of external space, just as proposed for auditory stimuli. However, another option is that remapping takes place within a “bodily map”, or, in other words, a spatial body representation (Longo et al., 2010; Medina & Coslett, 2010; Tamè et al., 2019). In this case, the perceived spatial properties of the body, such as its size, shape and posture, are adapted to the visual map.

Here, we separate these possibilities of spatial vs. bodily mapping by recalibrating stimulus positions on the hand; we used a ventriloquism aftereffect protocol that recalibrated stimulus positions at the fingertips and knuckles in outwards directions, with the origin of the outwards shift being the wrist. Two kinds of observations can differentiate between external and bodily maps in this paradigm.

First, if modulation of localization were related to external space, then localization should exhibit one common bias over all stimulus locations, and this common bias should reflect the average, outwardly directed offset across the trained stimulus locations. Notably, this average bias would also affect the perceived location of the wrist. In contrast, if modulation of localization were related to a change in perceived hand size, the bias at any given stimulus location should depend on the anatomical layout of a “scaled up” hand; thus, for example, the bias should be directed outwards for stimuli on the distal parts of the hand but remain stationary or point inwards for the proximal parts of the hand. Similarly, the perceived location of the wrist as the hand’s base should be perceived unchanged, as growth would appear to occur outwards, originating at the wrist.

Second, if localization changes were related to the change of a body map, then our recalibration procedure should affect not only direct localization but also other perceptual measures that rely on hand size estimations. For instance, when participants localize landmarks of the hand, such as the knuckles and fingertips (Longo & Haggard, 2010), they systematically underestimate hand length but overestimate hand width. The same systematic distortion is evident when participants localize tactile stimuli on the back of the hand (Mancini et al., 2011). Moreso, a related bias can affect spatial judgments that do not require explicit stimulus localization: Distance judgments between two simultaneously presented tactile stimuli, termed tactile distance judgments (TDJ), are larger for stimuli presented along the mediolateral (width) than along the proximodistal (length) axis of the hand’s back (Longo & Haggard, 2011; Miller et al., 2016). Thus, one and the same type of distortion reveals itself across multiple, conceptually different tasks and measures, supporting the idea that they all depend on one common, underlying representation of body (part) shape and size. This idea is further supported by the finding that both localization and TDJ are sensitive to experimentally induced changes of perceived body size. For instance, after using a hand-held tool for several minutes, participants localize tactile stimuli applied to the wrist and the elbow further apart than before tool use, suggesting that they now perceive their arm as elongated (Cardinali et al., 2009). The same manipulation also results in smaller TDJ on the forearm (Miller et al., 2014, 2017). Thus, these studies suggest that body representations expand when using tools that extend reaching space, and that the change of these representations then affects the different experimental measures we have discussed.

In the present study, we aim to establish the link between visuotactile ventriloquism aftereffects and presumed body representations, that is, underlying cognitive, map-like structures that underlie body processing. We imply an enlargement of hand size through visuotactile stimulation in which visual stimuli are presented further outwards than their tactile counterparts. We then test three aspects of visuotactile recalibration: First, we test whether localization by pointing to tactile-only targets reflects an enlargement of hand size as well. Second, we test whether the same recalibration procedure affects TDJ as another, independent measure, as parallel changes in the two independent tasks would indicate the involvement of a common, underlying body representation. Third, we assess explicit self-reports about perceived hand size to test the potential transfer of recalibration to an explicit body representation.

## Methods

### Participants

Twenty-five participants (16 females, aged 24.4 ± 4.8 years) took part in the study and were randomly assigned to groups receiving either constant or incremental recalibration (see next sections). To ensure that participants did not perform pointing movements with their non-dominant hand, we only included right-handed (n=22) or ambidextrous (n=3) participants, as assessed by the Edinburgh Handedness Inventory (Oldfield, 1971). We recruited participants with billboard advertisements posted at Bielefeld University and in digital platforms. They either received course credit or participated without compensation. All participants were free of sensory deficits, neurological, and psychological disorders by self-report. They were informed about the study’s procedures and gave written informed consent. The study was approved by the local ethics committee.

Due to a lack of prior studies about hand resizing using recalibration, we did not perform a formal power analysis. We decided on an a priori sample size of 25 participants, which we judged as the upper limit of feasibility, given that one assessment took about three hours and the study was performed during the COVID-19 pandemic. Our sample size is slightly above the range of other studies (10-20 participants) that have used similar procedures and shown either ventriloquism aftereffects (Bruns & Röder, 2019; Frissen et al., 2012) or hand anisotropies (Calzolari et al., 2017; Longo & Haggard, 2011; Miller et al., 2017).

### Procedures

The study consisted of two sessions, performed in randomized order. In one session, tactile and visual stimuli were presented in a spatially congruent manner; in the other session, they were not spatially congruent (congruent vs. incongruent, see next section). Each session started with a pre-test baseline assessment of localization-based hand maps and TDJ (see Figure 1). Then participants underwent a training phase that consisted of 15 iterations of visual-tactile stimulation, each followed by a localization task that assessed the potential recalibration of tactile localization. Subsequently, TDJ were assessed again, referred to as post-test. Finally, participants rated self-report items (see below). Participants did not move during the session, nor did they see their hand, so that visual and proprioceptive updating of hand position and size was not possible. One session lasted about 1-1.5 hours. Participants took a break between the two sessions.

**Figure 1:**
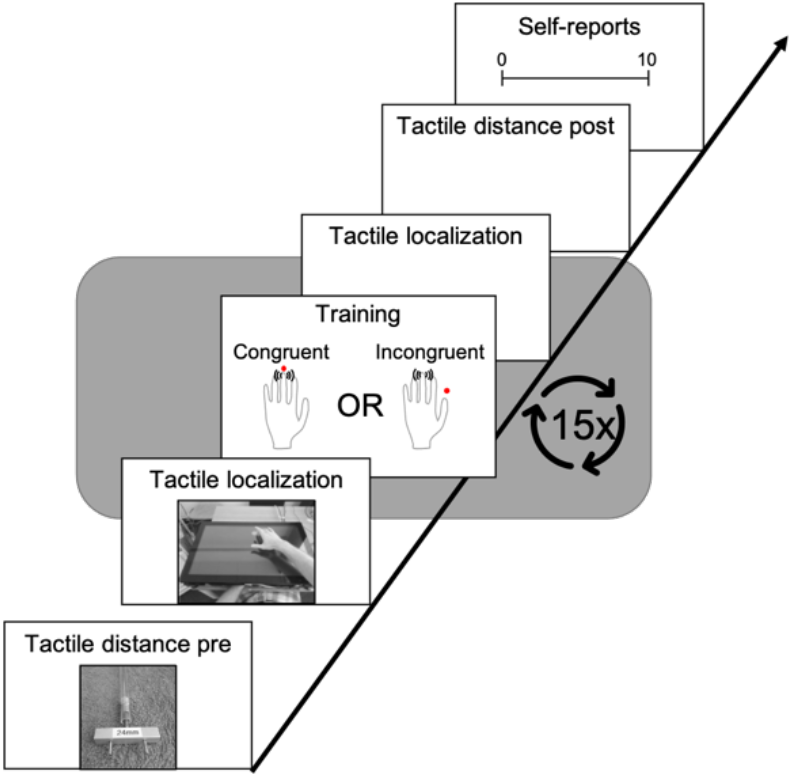
Study design. The study comprised a congruent and an incongruent recalibration session in randomized order. Each session consisted of a pre-test for and tactile localization-based hand maps and tactile distance judgments. Then the participants underwent 15 iterations, each comprising a recalibration phase and a tactile localization task. At the end of each session, the participants performed again the tactile distance task as a post-test and rated experiencing grown hand size and other potential perceptual changes (see text for details).

### Tactile localization and recalibration

Eight 1.8 cm diameter solenoid type tactile stimulators (Dancer Design, St. Helens, United Kingdom) were attached to participants’ palmar hand using adhesive rings. One stimulator was attached to the last segments of each digit, one each to the knuckles of D1 and D5, and one to the base of the hand, on the Carpal delta (see Figure 2A). The tactile stimulators were driven by a Neurocore (Neurocore, Hamburg, Germany), a standalone unit that can play and amplify stimulus waveforms with sub-millisecond precision upon receiving an external trigger. The stimuli were 200 Hz sine waves of 45 ms duration, presented at the maximum output intensity of the Neurocore. Time-precise triggering was achieved via the digital outputs of a Labjack U6 (Labjack, Lakewood, CO, United States) controlled via the LabJackPython library for the python programming language (www.python.org).

**Figure 2:**
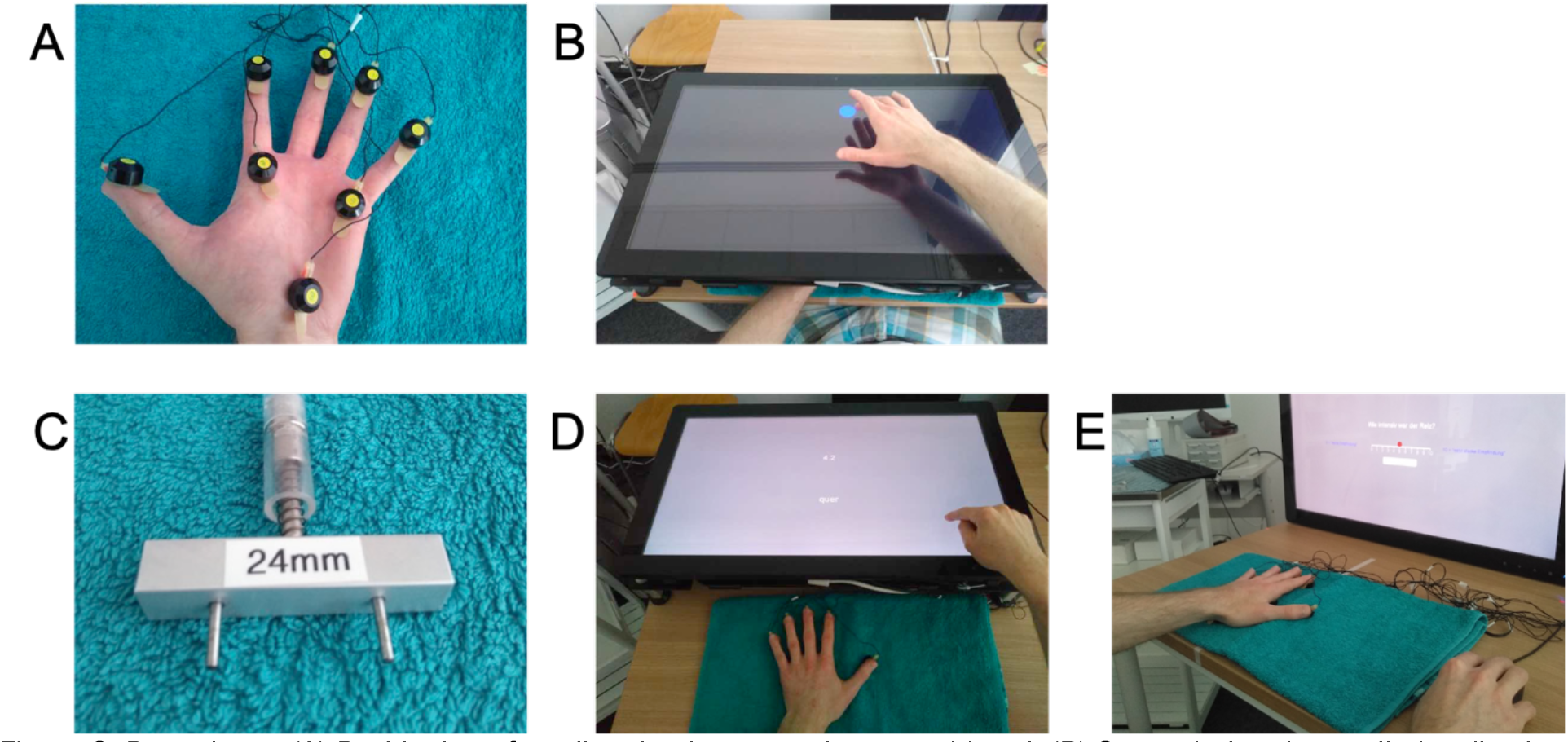
Procedures. (A) Positioning of tactile stimulators on the ventral hand. (B) Setup during the tactile localization phase. The participant’s hand was positioned palm-down on a towel underneath a mounted, horizontally placed touchscreen monitor. In each trial, the participant first touched a visually presented starting position (blue circle), then received a tactile stimulus in one of the positions indicated in A and then tapped the area of the touchscreen above the perceived stimulus location perpendicularly from above. (C) An exemplary stimulation device used for the assessment of tactile distance judgments. (D) Setup during the tactile distance judgment task. The participant closed the eyes and was then briefly touched with a stimulator like the one shown in C. Then the participant reproduced the perceived distance by sliding the right index finger across the surface of the touchscreen, reproducing a distance matching the length of the perceived stimulus distance. (E) Setup during the perceptual self-reports and the intensity ratings. The monitor was positioned vertically and the rating scales were operated using the mouse.

### Tactile localization and recalibration

The participant’s hand was placed palm down on the table surface. The towel under the hand prevented the sound of vibrotactile stimuli from being transduced to the table surface. Participants wore standard stereo headphones that played pink noise at an individually adjusted, comfortable volume to shield any potential, residual noise of the stimulators. A flatscreen monitor (iiyama ProLite T2753MSC, iiyama, Tokyo, Japan), mounted horizontally on an aluminum frame, was positioned over the participant’s hand. The touchscreen area of the monitor had a width of 60 and a height of 34 cm and was operated using a resolution of 1920 × 1080 px and a framerate of 60 Hz. The construction left sufficient space (6.9 cm) to place the hand underneath the construction in a comfortable position without touching any part of to the construction. The monitor’s touchscreen was elevated 12.6 cm above the table surface.

#### Tactile localization task

The localization task assessed localization-based hand maps (Longo & Haggard, 2010; Mancini et al., 2011; Medina & Duckett, 2017). In each trial, participants first saw a blue circle (3 cm diameter) on the touchscreen that served as a starting position for the participant’s localization movements. The starting positions were presented at a random position of the screen, leaving a margin of 15% of the width and height respectively at the sides. The randomization of starting locations was done to prevent stereotypical movements. The blue circle disappeared as soon as participants touched it and after a random time interval between 12 and 24 frames (200 - 400 ms), a tactile stimulus was presented randomly at one of the 8 hand positions. Participants then tapped the position on the touchscreen where they had perceived the stimulus. After releasing the finger from the touchscreen, the next trial started with the display of the next starting position. Each of the 8 stimulus positions was presented 10 times in the pre- and post-tests, and 5 times after each round of recalibration.

#### Recalibration task

The aim of the recalibration task was to test whether tactile position can be recalibrated via the ventriloquism aftereffect to induce an enlargement of represented hand size. We introduced spatial offsets between the positions of the tactile and the visual stimuli so that the visual stimuli corresponded to a hand map that was larger than the tactile hand map. It is debated whether recalibration works best in an incremental or in a constant way. During an incremental procedure, the offsets between tactile and visual stimuli are initially small and increase during the experiment. Whereas incremental prism adaptation has a stronger effect than constant adaptation on auditory recalibration in owls (Linkenhoker & Knudsen, 2002) and stronger motor adaptation aftereffects in humans (Kagerer et al., 1997), the constant method proved superior in a direct comparison of incremental and constant recalibration for the visual-auditory ventriloquism aftereffect (Bruns & Röder, 2019) and is common in both studies on visuotactile ventriloquism (after)effects (Samad & Shams, 2016, 2018) and on body illusions (Botvinick & Cohen, 1998). Due to these mixed prior reports, we used constant recalibration in half of our participants and incremental recalibration in the other half. Moreover, we included a factor *type of recalibration* (constant vs. incremental) in our analyses.

Recalibration used red circles with 1 cm diameter, presented for 3 frames (50 ms), presented in synchrony with the tactile stimuli. For each stimulus, one of the 8 vibrotactile stimulators was selected pseudo-randomly, with each stimulator repeated 25 times per iteration, resulting in 200 stimuli for one iteration (in total 3000 stimuli per session). Tactile-visual stimulus pairs were separated from the next pair by a variable interstimulus interval of 2-12 monitor frames (∼ 33-200 ms).

The position of the visual stimuli depended on the experimental condition (see Figure 3). In total, there were 4 recalibration conditions resulting from the combinations of the factors *congruency* (congruent vs. incongruent visuotactile stimulation) and *type of recalibration* (constant vs. incremental procedure). In all 4 conditions, visual stimuli were presented relative to the respective tactile locations reported by the individual participant in the pre-test. However, visual stimuli were then presented further outwards than the initial positions, with the intention to imply an enlargement of visual hand size compared to the hand size assessed via the tactile pre-test. In congruent resizing conditions, the mapping between tactile and visual stimulus positions was fixed. This mapping was anatomically congruent, which means that, for example, each time when the tip of D3 was stimulated tactually, the visual stimulus was presented in the location where the tip of D3 could be expected given that the hand size was scaled up (see next section). In contrast, this mapping was scrambled in incongruent conditions: each time a tactile stimulus was presented, any of the 7 other visual stimulus positions was selected randomly, so that tactile and visual stimulus locations did not match. In other words, in both congruent and incongruent conditions, participants were presented with visual stimuli that were located at comparable distances from the tactile stimuli and further outward. The only difference was whether they were anatomically congruent or not. Participants were instructed to pay attention to the red points that would appear on the screen as well as to the touch presented on their left hand.

**Figure 3:**
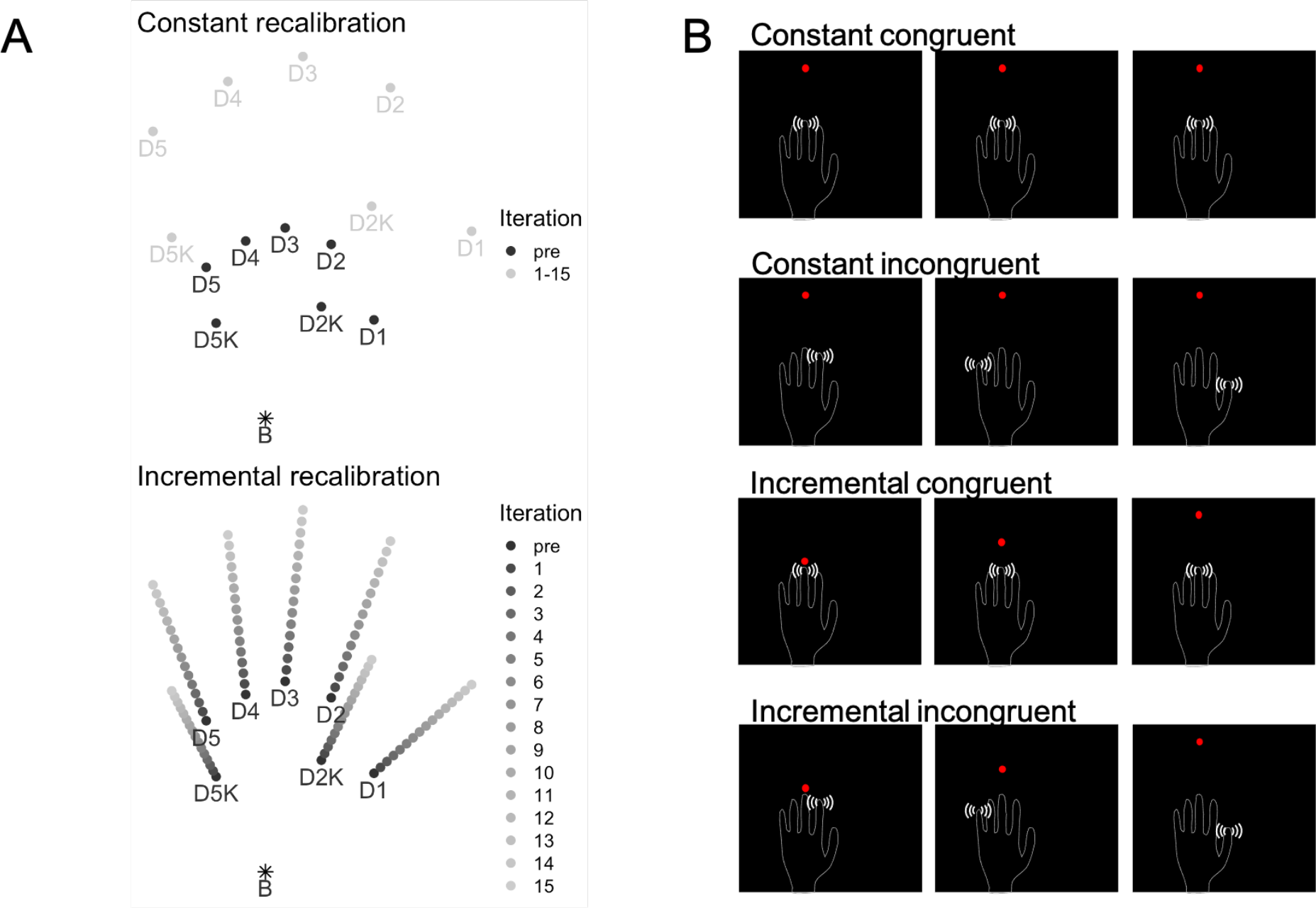
Experimental conditions. (A) The factor *type of recalibration* defined the development of spatial offset over 15 iterations of the task. In the constant recalibration, the offset was the same maximal offset over all iterations; in the incremental type, the offset increased gradually over the 15 iterations and reached the maximum offset in the last iteration. (B) The factor *congruency* defined the (anatomical) correspondence between the visual and the tactile stimuli. In the congruent condition, there was a fixed and anatomically congruent mapping between the stimuli whereas in the incongruent condition, the mapping was scrambled, that is, randomized over the stimulations. The two factors result in 4 types of conditions, depicted in B. The rows of images belong to the factor combinations; the three columns represent different iterations.

The scaling of the visual stimuli referred to the individual participant’s localization map, as assessed in the pre-test. We refer to hand size based on this map as the “original” size from here-on. We used median localization values to yield robust location estimates that are unaffected by potential outliers. We scaled the original hand map up with an affine transformation and then aligned the wrist/hand base position of the original and the transformed hand maps. Thus, the hand maps “grew out” of the hand base position at the end of the arm. Accordingly, visual stimuli were not offset at the base, i.e., the wrist stimulus. The maximum scaling factor was 1.9. In the incremental group, size increased in equally-sized steps with each of the 15 iterations. In the constant group, the maximum size was used during all 15 iterations (see Figure 3).

#### Practice block

For both the localization and recalibration task we familiarized participants with the procedures at the beginning of each session. The practice phase comprised two tactile localizations of each stimulator and a shorter version of the recalibration phase with 5 repetitions per stimulator and no spatial offsets.

### Tactile distance judgments

The aim of the TDJ task was to test whether the implied hand size change we attempted to induce with our recalibration procedure would affect subjective reports related to tactile-spatial perception that are not directly related to the stimuli employed to induce the perceived size change. We chose the TDJ task because previous research has indeed suggested that TDJ rely on an underlying hand size representation (Calzolari et al., 2017; Longo & Haggard, 2011; Miller et al., 2016).

In each trial, we touched the participant’s hand dorsum with a one of three stimulators (see Fig 2C) perpendicularly from above for about one second. Each of the three stimulators consisted of two small protruding metal rods interspaced by 2.4, 3.3, and 4.2 cm, respectively. A spring (force increase of 0.09 N/mm) attached to the pressure points standardized pressure application. Each stimulator was applied 14 times in both mediolateral and proximodistal direction. The order of stimulus size and direction was randomized. We instructed participants to pay attention to the stimulus and estimate the distance between the two rods of the stimulators. Directly following the stimulus, they reproduced the perceived distance by sliding their right hand’s index finger across the surface of a touchscreen monitor (Keizer et al., 2012). To prevent any bias resulting from (in)congruency between the direction of the stimuli and the direction of the reproduced distance, we divided the session into two parts in which the direction of the finger response movement on the screen was horizontal (left-right) and in the near-far direction, respectively. The order of these parts was randomized across participants. Participants kept their eyes closed during the entire procedure. Before data acquisition, participants briefly practiced the task.

### Perceptual self-reports

After the localization and TDJ task were complete, participants rated six statements that were presented in random order on the screen. They rated their agreement using an 11-point numerical scale that ranged from 0 (no agreement) to 10 (full agreement) and was operated with the computer mouse. The statements assessed whether, during the session, participants had perceived their hand to have become bigger or smaller; about how precisely they had perceived their hand’s location; and whether their perceived hand position had seemed to drift to the left or right (see Table 1). We expected statements about perceived size, but not precision or location, to be affected by recalibration.

**Table 1:** Perceptual self-reports assessed after recalibration

| The statements were: "during the course of the session I had increasingly the impression that..." |  |  |
| --- | --- | --- |
| # | wording | dimension |
| 1 | the perceived location of my left became more precise | Precision |
| 2 | the perceived location of my left became less precise | Precision |
| 3 | the perceived size of my left hand increased | Size |
| 4 | the perceived size of my left hand decreased | Size |
| 5 | the perceived location of my left hand drifted to the right | Location |
| 6 | the perceived location of my left hand drifted to the left | Location |

### Perceived Intensity

To ensure that participants had perceived all stimuli well, they rated the perceived intensities of the tactile stimuli using a numerical rating scale at the end of the session. The scale was presented on the screen and ranged from 0 (“no sensation”) to 10 (“very strong sensation”). Participants selected the corresponding value using the computer mouse (see Fig. 2E). The eight stimulators were presented in random order and each stimulator was presented 3 times. The results of the intensity ratings confirmed that stimuli were perceived with a moderate intensity and are presented in the supplementary material.

## Statistical Analysis

Data of one participant’s incongruent condition session was lost due to experimenter error. The remaining data of this participant were nonetheless used for analysis.

We assessed statistical significance of all results with linear mixed models (LMMs; Pinheiro & Bates, 2000). We initially included all relevant factors in any given LMM as fixed effects and included all main effects and 2-way interactions. We included 3-way interactions when they were of theoretical interest (see next sections).

As it is recommended to use a complex random effects structure (Barr et al., 2013), we initiated the models with a random intercept for participants and with all simple random slope effects and then iteratively reduced single random slope terms in case the model did not converge.

We followed up significant main effects or interactions with post-hoc tests based on estimated marginal means. We corrected multiple tests with the false discovery rate (FDR; Benjamini & Hochberg, 1995).

### Tactile localization maps

The aim of the analysis was to detect potential changes in implicit hand size, as expressed by the tactile localization task, over the sessions. We hypothesized that the congruent mapping of visual and tactile stimuli during the size recalibration procedure would change the size of a presumed hand representation more than the incongruent mapping. A few localization responses were very far away from the localization centroids (average position for stimulation sites), presumably due to accidentally touching the monitor during the reaching movement. We discarded localizations which were more than 20 cm from the respective centroid, which was the case in 32 localizations (<0.01% of the data).

We addressed three questions in separate analyses: first, whether congruent recalibration enlarged perceived hand size overall from before to after the recalibration procedure; second, the evolution of change across the 15 iterations of our recalibration procedure; and third, location-specific effects. We describe these three analyses here in turn:

#### (1) Overall size enlargement from before to after recalibration

We computed the localization centroids for the hand base position (point “B” in Figure 3A) as the average for the pretest as well as separately for each recalibration iteration. As dependent measure, we then computed the Euclidian distance of each localization from its iteration’s hand base centroid. These distances express how far each stimulus was localized from the hand base, and this distance should get larger when hand size is perceived as enlarged. Given that the hand base centroid was computed for each iteration, distance values are internally corrected for potential drifts of perceived hand position (including the base). For the analysis of hand size change, we ran two LMMs. First, we performed a pre-post comparison coded as a within-subject fixed factor *time* (levels: pre, post). As other fixed factors, we included the within-subject factor *congruency* (levels: congruent, incongruent session) and the between-subject factor *type of recalibration* (levels: constant, incremental). The hypothesis of a size change over time following congruent recalibration is expressed by the *congruency* × *time* interaction. Because the LMM predicts the average of the distance values, we refer to the estimated marginal mean derived from the LMM as the *hand size index* in the following sections. To allow testing whether *type of recalibration* influences the hand size index, we included the *congruency* × *time* × *type of recalibration* interaction.

#### (2) Evolution of perceived size change over time

We analyzed how the hand size index evolved over the 15 iterations of the recalibration procedure. We predicted a steeper size change slope for congruent than for incongruent recalibration. We included the continuous predictor *iteration* (pre-test, coded as iteration zero, and the following the 15 iterations), as well as *congruency*. To follow up significant effects of *iteration*, or interactions involving *iteration*, we performed a trend analysis, that is, statistical comparison of regression coefficients between the levels of the involved factors.

#### (3) Landmark- and direction-specific analysis

We analyzed changes in localization in detail by comparing pre-test vs. the post-test localization of the 8 hand landmarks separately and analyzed their recalibration in the near-far vs. the left-right direction. Recall that the pattern of recalibration of the different landmarks is indicative of whether the recalibration procedure altered a representation of hand size or whether all stimuli are recalibrated according to one common average independent of any underlying hand representation. Visual stimuli were offset outwards, i.e., continuing the along the line connecting the wrist with the respective landmark (see Figure 3). Hand map-based recalibration should, accordingly, result in larger adjustments for more distal landmarks. Similarly, left landmarks (e.g., D5) should recalibrate to the left and right landmarks (e.g., D1) to the right, producing a “fan-shaped” recalibration pattern. In contrast, stereotypical or average recalibration for all hand landmarks would result if no hand size representation were involved. We calculated change scores for the near-far and the left-right localization change by subtracting the average localization value of the pre-test from the localizations of the post-test. Positive values indicate a change in the far or the right direction, respectively. We then computed two LMMs, one for the near-far, and one for the left-right localization change scores with fixed factors *congruency* (levels: congruent, incongruent session), and *hand landmark* (levels: B, D1, D2, D3, D4, D5, D1K, D5K).

### Tactile distance judgments

The aim was to detect whether TDJ changed from pre- to post-test, and whether a potential change may depend on visual-tactile stimulation being congruent during recalibration. The LMM included fixed factors *congruency* and *time*. We included *direction* (mediolateral vs. proximodistal) to test for the typically reported anisotropies of width and length. Moreover, we included distance (2.4, 3.3, 4.2 cm) to test whether larger distances were judged larger. The hypothesized change of perceived stimulus size following congruent (but not incongruent) recalibration is expressed by the *congruency* × *time* interaction.

### Perceptual self-reports

We analyzed self-reports with an LMM that included the fixed factors *congruency* and *question* (levels: questions 1-6, see above).

### Used software and availability of data and code

All statistics were performed with the R statistical computing language (v4.2.1, https://cran.r-project.org) in the RStudio environment (v2021.09.1, www.rstudio.com). We fitted and analyzed LMMs with the *lme4* (Bates et al., 2014) and *lmerTest* packages (Kuznetsova et al., 2014) and performed post-hoc comparisons and trend analyses with the *emmeans* package (Lenth et al., 2019). We created figures with the *ggplot2* package (Wickham, 2009).

All data and code to reproduce the statistical results and figures presented in this article are available on the website of the Open Science Framework under https://osf.io/bqpy2/.

## Results

### Tactile localization maps

The change of the localization map is shown in Figure 4A, which gives an impression of both magnitude and direction of recalibration.

**Figure 4:**
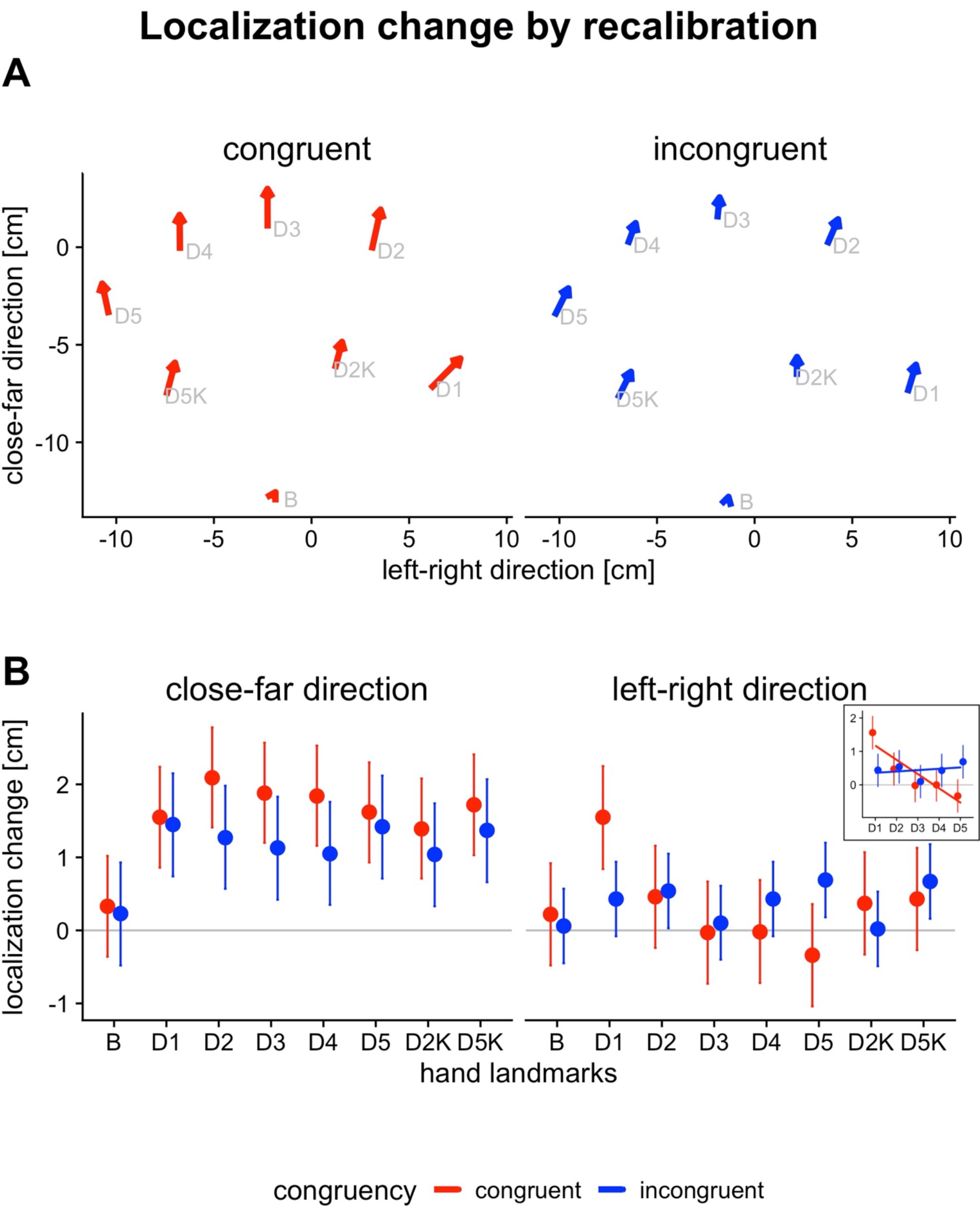
Change in localization-based hand maps by visuotactile recalibration. (A) Hand localization maps before and after recalibration. The localization map at pre-test is indicated by the starting point and the post-test by the tip of the arrows. The arrows’ length therefore indicates the magnitude and their inclination the direction of recalibration. The units are cm; the origin (0,0) is the middle of the touchscreen monitor. (B) Localization change (before vs. after recalibration) in the near-far and left-right direction. The points represent estimated marginal means derived from the linear mixed models with 95% confidence intervals as error bars. For the near-far direction, there was a significant change for all body sites except for the hand base (grey letter “B”), for which no visuotactile offsets were used. The left-to-right change depended on congruency and the hand landmarks. Notably, D1 more strongly recalibrated to the left and D5 more strongly recalibrated to the right in the congruent compared to the incongruent condition. The subplot in the upper-right corner of panel B presents the result of an additional submodel, computed post-hoc, that includes only the fingertip landmarks (D1 to D5). A significantly negative regression slope was evident only for congruent recalibration, indicating that these landmarks resembled an expansion of hand size (see text for details). The result pattern is consistent with an outwards-directed, “fan-shaped” enlargement of the hand map that follows the direction of visuotactile offsets during the recalibration task (compare Figure 3A).

### Size increase of the localization map

*Pre-post comparison*. The size of localization-based hand maps increased from before to after recalibration, especially in the congruent condition (see Figure 5A). The greater size change in the congruent than in the incongruent condition was evident in a significant *congruency* × *time* interaction (F(1, 6586) = 17.53, *p* < 0.001). Post-hoc comparisons indicated that perceived hand size was significantly larger after than before recalibration for both congruent (difference_pre - post_ = -1.58 cm, t(28.96) = -5.26, *p* < 0.001) and incongruent conditions (difference_pre - post_ = -0.77 cm, t(29.09) = -2.55, *p* = 0.016), with a larger difference between pre and post (effect of *time*) in the congruent than the incongruent condition. The interaction between *time* and *type of recalibration* was not significant (F(1, 23) = 0.06, *p* = 0.814) indicating that both recalibration methods, incremental and constant, were similarly effective. There were no other significant terms in the LMM (see Table 2).

**Table 2:**
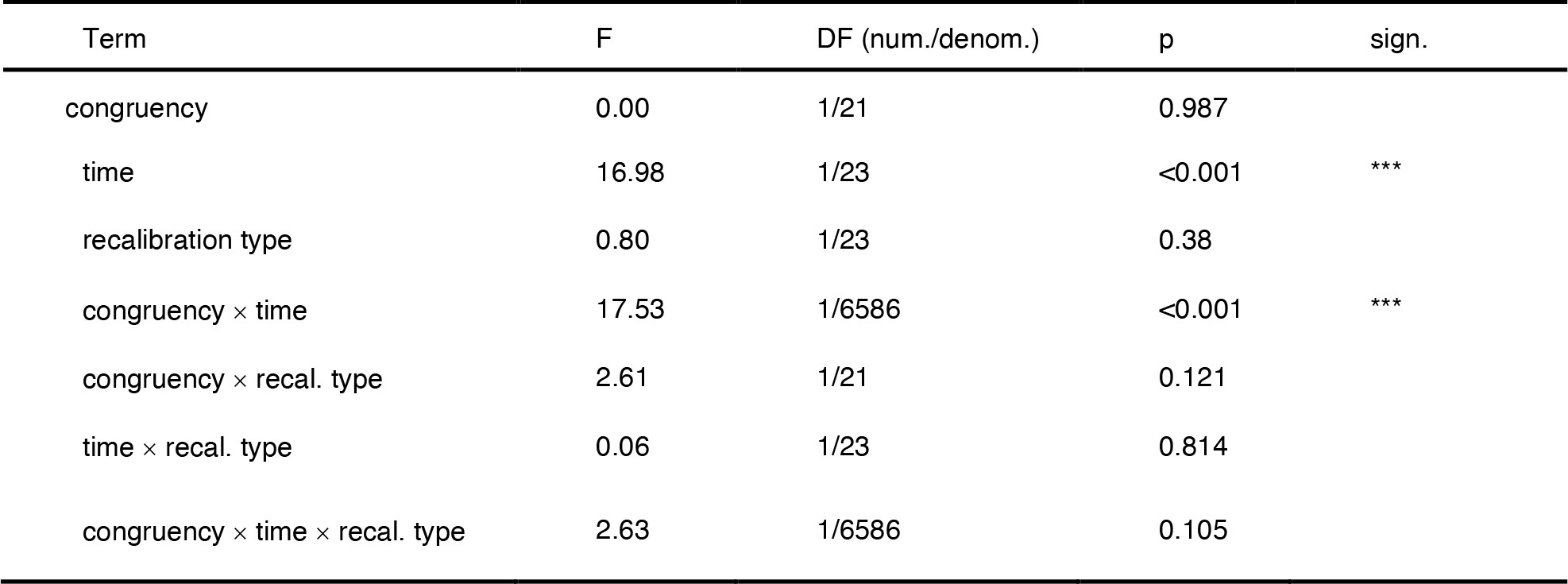
Results (ANOVA table) from the linear mixed model comparing hand size before and after recalibration.

**Figure 5:**
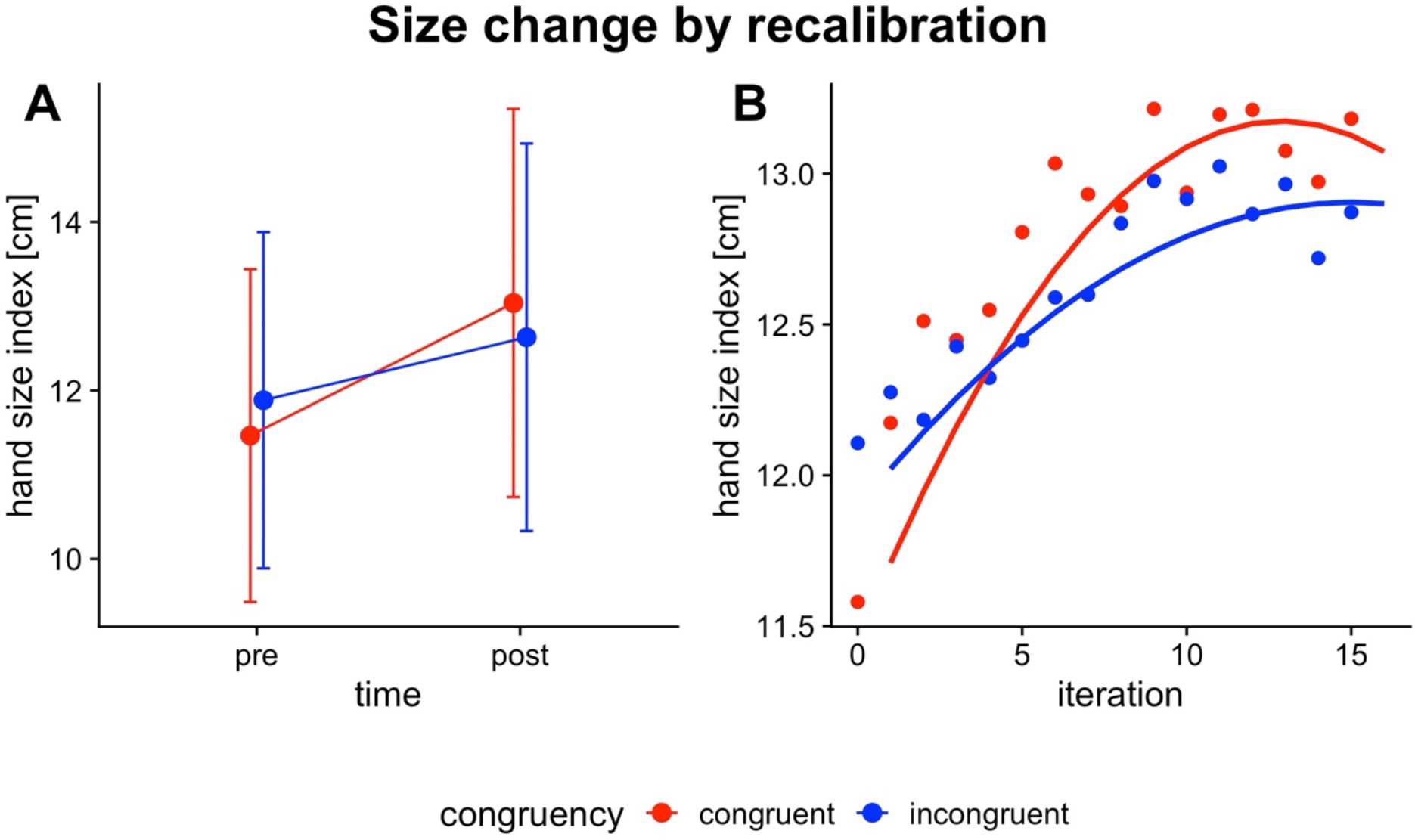
Change in the size of localization-based hand maps by visuotactile recalibration. (A) Interaction plot for *time* x *congruency*. The points are estimated marginal means derived from the linear mixed model (LMM) and the error bars are 95% confidence intervals. (B) Regression curves derived from linear and quadratic components of the LMM that assessed hand size change over the pre-test (iteration zero) and the following 15 recalibration iterations (see text for details). The hand size index in cm units (y-axis) was computed as the average distance of localizations from the hand base (letter “B” Figure 4A).

*Development over time*. The LMM over the 15 recalibration iterations fitted both a linear and a quadratic hand size change component. The linear component rose significantly steeper for the congruent compared to the incongruent condition, expressed as an *iteration* x *congruency* interaction in the LMM (F(1, 25661) = 12.59, *p* = 0.001). A corresponding interaction was found for the quadratic component: it was significantly more negative (i.e., decelerated more strongly) for the congruent compared to the incongruent condition (*iteration*_*squared*_ x *congruency* interaction: F(1, 24156) = 7.86, *p* = 0.005). As illustrated in Fig 5B, recalibration increased strongly in early iterations of the congruent condition and then reached a plateau. This progression implies fast adaptation early on during the recalibration phase and was less salient in the incongruent condition, as implied by the statistical results. Coefficients for the linear and quadratic components are presented in the supplementary material.

### Direction of recalibration

For both the close-far and the left-right direction, there were significant interactions between *congruency* and *hand landmark* (close-far direction: F(7, 3776) = 2.82, *p* = 0.006; left-right direction: F(7, 3776) = 9.27, *p* < 0.001) indicating that the magnitude of recalibration depended on congruency, but this effect differed between the 8 hand landmarks.

For both congruency conditions all hand landmarks significantly recalibrated in the far direction, except for the hand base. At the hand base (where visuo-tactile stimuli were presented without offset) the change did not significantly differ from zero in both congruency conditions (see Fig 4B, left side). The magnitude of recalibration in the far direction was significantly stronger in the congruent than the incongruent condition for the tip of D2 (0.82 cm, t(35.72) = 2.04, p = 0.048; FDR-corrected), and marginally significant for the tips of D3 (0.75 cm, t(35.72) = 1.88, p = 0.068), and D4 (0.79 cm, t(35.72) = 1.96, p = 0.057).

For the left-right direction, there was a significant rightward recalibration in the congruent condition for D1 (1.55 cm, t(32.3) = 4.49, p = 0.001; see Figure 4B, right side). Congruent and incongruent recalibration differed significantly for two landmarks: At D1, rightward recalibration was significantly stronger for congruent than incongruent stimulation (difference_congr. – incongr._: 1.1 cm, t(40.4) = 2.79, p = 0.008). Similarly, for D5, recalibration was leftwards for congruent and rightwards for incongruent stimulation (difference_congr. – incongr._: -1 cm, t(40.3) = 2.58, p = 0.013). This recalibration pattern is well in line with an up-scaling of hand size, with lateral landmarks moving outward, but central landmarks remaining near their original location with respect to left and right. We ran an additional LMM post-hoc to directly test the idea that recalibration was systematically related to landmark position. It included the left-right position of the five fingertip landmarks (D1 to D5) and treated these positions as a continuous predictor to fit linear regression slopes. The regression slope was significantly negative for congruent recalibration (*b* = -0.42, t(2349) = -8.87, p < 0.001) but non-significant and parallel to the x-axis for the incongruent recalibration (*b* = 0.04, t(2349) = 0.84, p = 0.399); moreover, the slopes of the two recalibration conditions were significantly different (difference_congr. – incongr._: -0.47, t(2349) = -6.87, p < 0.001). Thus, landmark-specific recalibration consistent with perceived hand size enlargement was exclusive to congruent recalibration.

### Tactile distance judgments

Next, we analyzed whether recalibration affected TDJ – a result that would support the existence of an underlying hand size representation.

Participants judged larger stimulus sizes as larger (main effect of *size*: F(2, 24) = 139.6, *p* < 0.001; see Figure 6); this result serves as a manipulation check, confirming that participants differentiated the different distances in our experiment. Tactile distances were judged significantly larger when the stimuli were applied in the mediolateral as compared to the proximodistal direction (main effect of *direction*: F(1, 24) = 65.07, *p* < 0.001) and confirms the common finding of width-length anisotropies in TDJ on the hand dorsum (see Figure 6). The results are presented in detail in the supplementary material.

**Figure 6:**
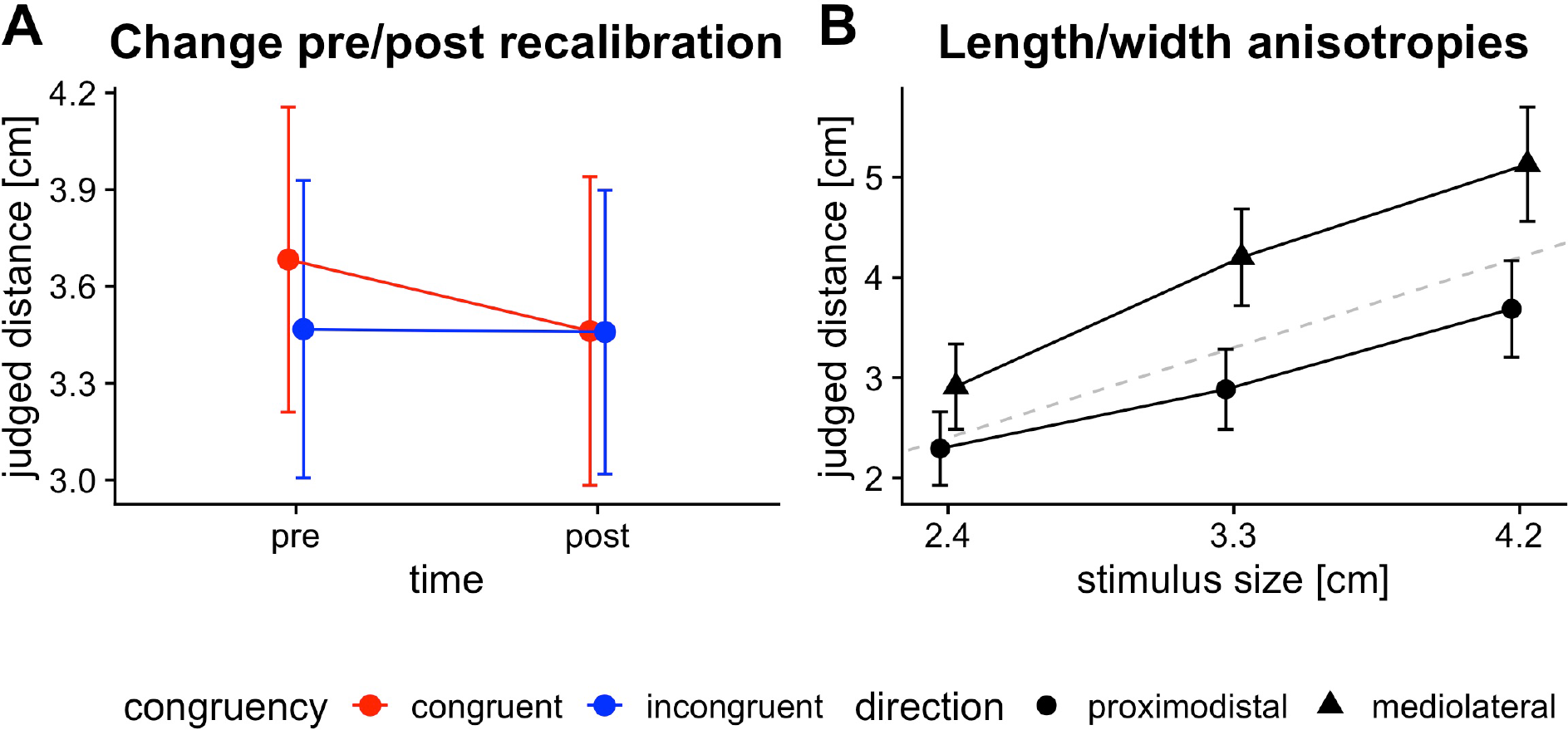
Change in tactile distance judgments (TDJ) by recalibration and length/width anisotropies. (A) Results from the linear mixed model comparing the TDJ before and after congruent and incongruent recalibration. The points are estimated marginal means (EMMs) from the model with 95% confidence intervals as error bars. The plot highlights a statistically significant interaction between *congruency* and *time* (pre vs. post) (see text for details). (B) EMM and 95% confidence intervals of length vs. width anisotropies in TDJ. The dotted line indicates perfect correspondence between actual size (x-axis) and judged size (y-axis). As has been commonly reported, distance stimuli in mediolateral direction were over- and stimuli in proximodistal direction underestimated.

### Size change by recalibration

TDJ decreased from before to after recalibration, but only in the congruent, evident in a *congruency* × *time* interaction (F(1, 7894) = 16.95, *p* < 0.001; see Fig 6A and Table 3), combined with post-hoc tests that were significant for the congruent but not the incongruent condition (see Table 4).

**Table 3:**
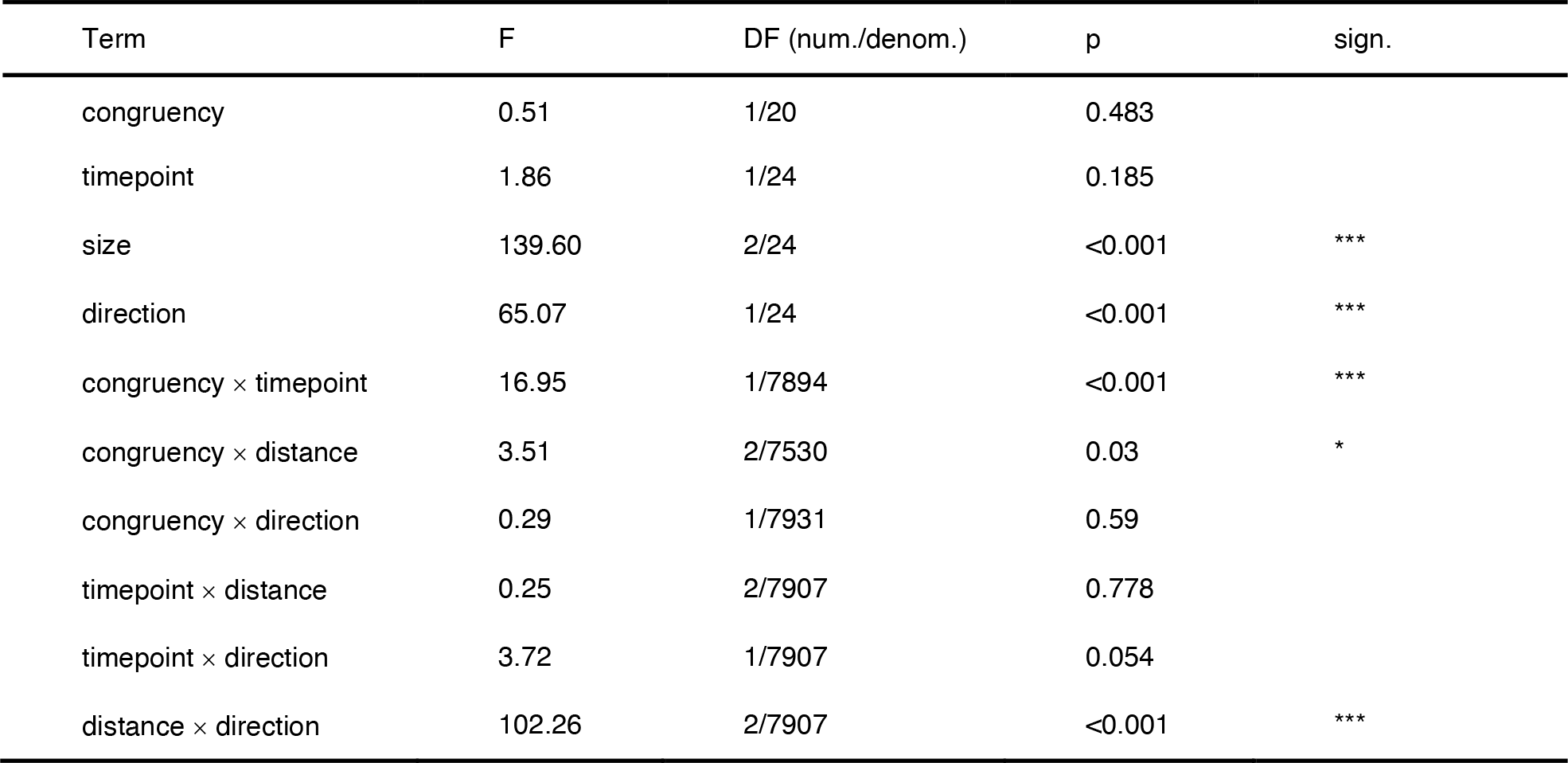
Results (ANOVA table) from the linear mixed model comparing tactile distance judgments beofore and after recalibration.

**Table 4:** Post-hoc comparisons between tactile distance judgements before and after congruent and incongruent recalibration.

| congruency | contrast | estimate | SE | DF | t ratio | lower CL | upper CL | p | sign. |
| --- | --- | --- | --- | --- | --- | --- | --- | --- | --- |
| congruent | pre - post | 0.22 | 0.09 | 28.63 | 2.51 | 0.04 | 0.40 | 0.018 | * |
| incongruent | pre - post | 0.01 | 0.09 | 28.63 | 0.10 | -0.17 | 0.19 | 0.924 |  |

### Perceptual self-reports

Recalibration congruency did not affect the responses to the perceptual self-report items (see Table 6 for the absence of main effects and interactions involving the fixed factor *congruency*).

**Table 6:**
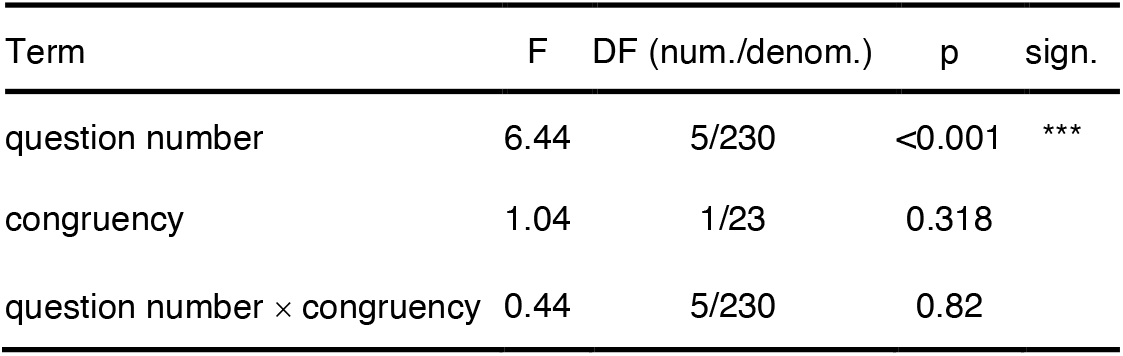
Results (ANOVA table) from the linear mixed model that compared the perceptual self-reports between congruent and incongruent recalibration.

Irrespective of the manipulation, the level of agreement differed across the six questions (main effect of *question*: F(5,230) = 6.44, *p* < 0.001; see Fig 7). For example, irrespective of *congruency*, participants more strongly agreed with the statement that their hand size had increased rather than decreased (difference_Q3 – Q4_ = 1.69, t(230)=3.12, *p* = 0.007), drifted right (difference_Q3 – Q5_ = 2.38, t(230)=4.39, *p* < 0.001), or drifted left (difference_Q3 – Q6_ = 1.35, t(230)=2.91, *p* = 0.031). However, participants did not more strongly agree that their hand size had increased than with it having become more or less precise.

**Figure 7:**
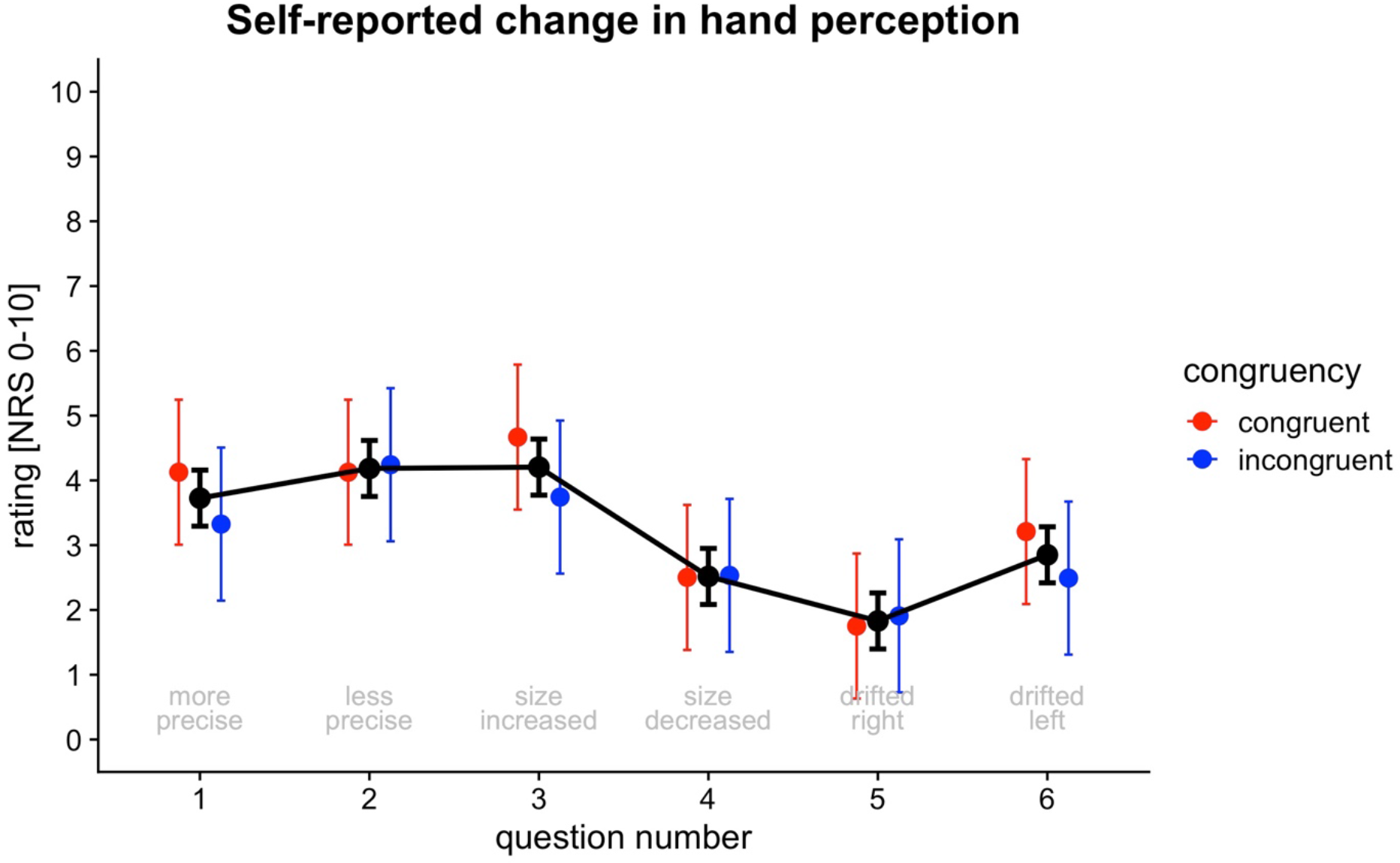
Self-reports on change in hand perception assessed after recalibration. The figure shows results from the linear mixed model that compares the ratings for different questions between the congruent and incongruent recalibration conditions. Points are estimated marginal means (EMMs) with 95% confidence intervals as error bars. The colored symbols represent the EMMs for the conditions separately. The black symbols for combined EMMs corresponding to the main effect of *question*. Separate ratings for congruent vs. incongruent recalibration are shown for descriptive reasons, but did were not statistically different from each other for any of the questions (see text for details).

## Discussion

In this study, we presented visual stimuli paired with tactile stimuli to a hand, with the visual stimuli implying an enlarged hand size. Our procedure is based on the visuotactile ventriloquism aftereffect that evokes recalibration of tactile stimuli. We used a congruent recalibration condition with a fixed mapping between visual and tactile stimuli and, as a control procedure, an incongruent condition where the mapping was scrambled. We tested whether these visuotactile sensory recalibration procedures differentially affected localization of unimodal tactile stimuli reported by hand pointing, TDJ, and the explicitly reported experience of a change in hand size. We report three main findings: (1) Following congruent recalibration, participants localized unimodal tactile stimuli like they aimed at larger hands than before recalibration or after incongruent recalibration. The direction of recalibration across the eight tested landmarks on the hand was consistent with the interpretation that recalibration was based on the size change of a hand map that had grown in an anatomically plausible way, as if the map were “scaled up”. (2) Tactile distance stimuli in the TDJ task were perceived as shorter following congruent recalibration, indicating effects of recalibration on another, independent measure of hand size representation. (3) Perceptual self-reports about consciously perceived hand size increase were not affected by stimulus congruency during recalibration. Thus, our recalibration congruency manipulation influenced two independent behavioral assessments, but not conscious self-reports. This result pattern suggests that congruent recalibration influenced an implicit size representation of the left hand. In addition, our results imply that the visuotactile ventriloquism aftereffect takes place within a plastic body map that adapts to sensory recalibration. Last but not least, our study shows that simple, bimodal stimuli without any body-related semantics are sufficient to change body representations, which extends our understanding about the minimal requirements necessary to influence body perception using multisensory stimuli. We will briefly discuss these topics in the following sections.

### Size increase of localization-based hand maps

With the tactile localization task, we tested whether the visually induced outwards recalibration of touch results in systematic localization bias also when tactile stimuli are no longer accompanied by visual stimuli. Such an effect would imply that the spatial map employed to execute reaches to tactually indicated body locations has increased in size. Participants’ pointing responses were indeed consistent with a presumed enlargement of their hand map. Although this effect was observable both after spatially congruent and incongruent recalibration, it was stronger in the congruent case, suggesting a specific effect of anatomically congruent visuotactile recalibration on the hand map employed for unisensory, tactile localization. In other words, modulation of hand size rested on specific, meaningful pairings of tactile and visual stimulus locations and was not achievable by visual stimuli simply occurring within a larger area of space while touch occurred on the hand.

We report evidence that participants in fact aimed their pointing movements towards a hand size representation that had been “scaled up” in an anatomically plausible way. First, we observed an increase in a hand size index that we computed by averaging distances between the wrist position and other hand landmarks. Specifically, the increase in the hand size index suggests that the perceived distance between the wrist and other landmarks increased, indicating that landmarks were remapped relative to each other, and the resulting hand map covered a larger portion of space following recalibration (cf. Cardinali et al., 2009). Second, analysis of recalibration for individual hand landmarks in the close-far and the left-right directions further supports this conclusion. In both the congruent and the incongruent condition, the wrist landmark, at which no recalibration was applied, remained stationary over the sessions whereas the other landmarks were recalibrated in the distal direction. The interpretation that the hand is “scaled up” gets further support from the different effects of congruent and incongruent recalibration. “Scaling up” in an anatomically plausible way should express itself in a “fan-shaped” pattern of recalibration: The fingertips should recalibrate in the distal direction; D1 and D5 as the inner and outer landmarks should also recalibrate to the right and to the left, respectively. We observed this predicted localization pattern with congruent, but not with incongruent visuotactile recalibration. The association between the digit number and the right-left direction of recalibration was also evident when analyzing the gradient across all five fingertip landmarks together: the hand landmarks on the right side recalibrated to the right, the ones on the left to the left and the intermediate ones stayed stationary. For incongruent recalibration the hand-shape pattern was absent; instead, all fingertip positions recalibrated towards the right, presumably in the direction of the body midline (Fuchs et al., 2016; Wann & Ibrahim, 1992).

Third, the progress of resizing over 15 iterations of recalibration was reminiscent of a learning curve that steeply rises and then flattens out (compare Fig 4C, congruent condition). This learning-like change was prominent in the congruent, but less so in the incongruent mapping. This result suggests that our cognitive system gradually relearns the bodily dimensions.

Based on these three aspects of our results, we argue that the interpretation that is most consistent with our present findings is that an underlying hand size representation was increased by, specifically, the congruent recalibration procedure and that this representation affected behavior in the localization task.

One might, instead, propose that our experimental results may stem from motor adaptation. In this view, participants merely adapted their pointing behavior to reach further while perceiving unisensory tactile stimuli at their unchanged spatial locations even after the ventriloquism procedure. This interpretation is unlikely for at least three reasons. First, if participants’ behavior were due to motor adaptation, one would expect a similar displacement for all tactile locations including the hand base location, unless one wants to assume stimulus-specific adaptation. It has been shown that reaches to two targets can be adapted independently from each other (Hirashima & Nozaki, 2012).

However, whether this concept can be extended to eight stimulus positions at once is unknown and at this point a rather far-fetched explanation for our results. Second, the ventriloquism protocol is designed to avoid motor adaptation in that participants never perform pointing movements to the bimodal, visuotactile stimuli during the recalibration procedure (see Samad & Shams, 2018 who applied the same logic to rule out motor adaptation). Thus, one would have to assume that purely observing spatially disparate multisensory stimuli results in motor adaptation to the stimuli’s unisensory constituents. We are unaware of any evidence for such motor-adaptive mechanisms. Third, motor adaptation cannot explain why distance was judged differently after our recalibration procedure, even more as the TDJ task involved stimulus locations that were never used during recalibration.

Finally, our study sheds some light on the effectiveness of constant vs. incremental visuotactile ventriloquism protocols. A previous study found that constant recalibration was superior for the audiovisual ventriloquism aftereffect (Bruns & Röder, 2019). In our study, the protocols lead to comparable effects and we therefore did not replicate the superiority of the constant protocol for the visuotactile case. This indicates that the effectiveness of protocols might depend on the sensory modalities, in line with the observation that incremental protocols have proved superior for motor adaptation (Kagerer et al., 1997).

### Decreased tactile distance judgments following recalibration

A more stringent test for the involvement of an underlying body representation in visuotactile recalibration is to test for modulation of performance in a task that is independent of the particular stimulus type and locations used in the recalibration procedure. We addressed this requirement with the TDJ task, which involved stimuli that were not used for recalibration – both regarding their location and quality – and required stimulus localization only implicitly, as participants had to derive the distance between two stimuli rather than explicitly pointing to them.

From before to after recalibration, the TDJ decreased, that is, distances were judged shorter after than before recalibration. Importantly, the TDJ decrease was stronger following congruent than incongruent recalibration, suggesting that it was related to our recalibration manipulation rather than adaptation effects which can occur in the TDJ task when stimuli are applied repeatedly (Calzolari et al., 2017).

It might strike as surprising that TDJ became smaller rather than larger with implied hand growth. However, the direction of the relationship between size representation and TDJ appears to be paradigm-specific. One way to conceive TDJ is that the size of the stimulus is judged relative to the internal representation of the size of the body part (cf. Tosi et al., 2022). For example, if the same distance stimulus is presented on a hand that is represented as relatively large, the stimulus should be judged as relatively large, too, because the two tactually stimulated locations that indicate the to-be-judged distance should be further apart in space if they are located on the same landmarks of a larger hand. Some studies using manipulations of body representations are in line with this view. For example, TDJ on the finger increase when perceived finger length is extended via a proprioceptive illusion involving tendon vibration (de Vignemont et al., 2005). However, TDJ along the arm usually decrease when a tool that extends the reaching space is used over a period of time (Miller et al., 2014, 2017). This tool use manipulation is otherwise associated with effects that imply that tool use extends perceived arm length: Following tool use, pointing movements towards body landmarks, for example towards the wrist and the elbow, change such that the two landmarks are localized further apart, implying an increased forearm size. Moreover, the kinematic characteristics of arm movements after tool use resemble those that would be expected if the arm were longer than it actually is (Cardinali et al., 2009). A conceptual framework that aligns the diverging findings about the relationship between size representation and TDJ is, so far, lacking (but see Miller et al., 2014 for a discussion of competing theoretical accounts). Importantly, TDJ decrease in tool use experiments has been reported consistently, and our TDJ pattern is comparable to these reports.

In sum, participants’ behavior in tactile length judgments was affected by recalibration. The decrease of TDJ in the context of an enlarged hand representation is well in line with an equivalent effect in the context of enlarged arm representation after tool use. Thus, our recalibration procedure affected both tactile localization and distance judgments in a coherent manner, supporting the conclusion that recalibration affected a common underlying representation of hand size.

### Effects of congruency did not transfer to perceptual self-reports

Participants, on average, agreed more strongly with the statement that their hand had perceptually grown than that it had shrunk. However, the ratings were not influenced by the experimental congruency manipulation. This result is in line with stimuli being consistently presented with an outwards-offset from the hand and participants also indicating an enlarged hand in the localization task in both congruency conditions. However, the localization map was significantly larger and TDJ shorter following congruent than incongruent recalibration and this effect was not present in the self-reports. This indicates that at least the effect of congruency on the perceptual hand size representation was implicit, because it was observable behaviorally in localization and TDJ but absent in (explicit) self-reports.

### Visuotactile recalibration updates an underlying spatial body representation

When being exposed to sensory information or sensorimotor contingencies that imply a change of spatial bodily features (like position, shape, or size), cognitive processes update to incorporate these contingencies. Such updating is evident, for instance, during the extension of grasping space when using tools (Cardinali et al., 2009; Miller et al., 2014, 2017) and visual information implying touch located on an artificial hand (Botvinick & Cohen, 1998). Visuotactile stimulation implying an enlarged hand, as employed here, is another driver for such updating. The underlying, updated cognitive architecture is often termed a “representation”, because changes are not restricted to the particular instances that participants experienced – which could be explained by simple associations –, but also to stimuli that have not been experienced with the updated contingency. Moreover, the concept of “representation” implies the use of an underlying structure or process that is not merely inherent to one given task, but underlies multiple tasks, optimally both with respect to the sampled sensory quality as well as the mode of reporting. In the present study, we utilized a multi-task approach to test whether a “representation” is affected by using two independent behavioral tasks that might be guided by an implicit hand size representation: localization-based hand maps and TDJ. TDJ are independent from the localization task both with respect to the employed stimuli (brief vibratory stimuli vs. a distance stimulator applied manually on the hand dorsum where no other stimuli had been applied) and the judged attribute of the stimulus (location vs. distance). Hence, any transfer to the TDJ cannot be based on simple associations. In the present study, we indeed found effects across the two independent behavioral tasks. Both the pattern of recalibration across the hand’s landmarks for tactile localization and TDJ were modulated by recalibration congruency in a manner that is consistent with a common, underlying, “scaled up” hand map. In contrast, the result pattern across tasks is neither consistent with simple alignment of touch with visual stimulation nor motor adaptation as drivers of the observed behavior.

We interpret the enlargement of a hand size representation as an implicit effect. Both tasks, localization and TDJ, assess perceived hand size indirectly via biases in the observed behavior. In the localization task, our estimate of represented hand size emerges from the reported locations of hand landmarks relative to each other. In TDJ, no size estimate is derived, but changes in represented hand size are inferred from distance judgments of diverse locations on the hand. Participants are probably unaware of changes in their behavior and the effect on the size metric that is derived from it. This conjecture is in line with the results of our questionnaire, which did not reflect the effects of congruency of our behavioral tasks.

Even if the present results favor the existence of a common underlying representation, the two tasks employ vastly different measures, prohibiting the conclusion that they are both identically affected by recalibration. Some studies have tested for a common, underlying representation across tasks by correlating performance between them; these correlations were surprisingly low. One study, for example, assessed hand size anisotropies using a landmark localization task and TDJ but did not observe a significant correlation between the tasks (Longo & Morcom, 2016). Another study (Matsumiya, 2022) assessed distortions of localization-based hand maps using both finger pointing and eye movements and found weaker distortions for eye movements, suggesting that hand shape representations may be effector-specific. Such lack of strong relationships may imply differences in the underlying body representations, but may also be related to other task differences. Here, we did not further explore the relationship between tactile localization and TDJ because the N of 25 is considered too low for meaningful correlation analysis (Schönbrodt & Perugini, 2013).

### Update of body size representation is based on simple visuotactile processes and works without body semantics

Our study has implications for the conceptualization of both visuotactile ventriloquism aftereffects and body representations. Many experiments that have aimed to manipulate body representations have used methods of prolonged sensory conflict that invoke sensory recalibration (Bellan et al., 2015; Botvinick & Cohen, 1998; Holmes et al., 2004; Kilteni et al., 2012; Lackner, 1988) but rarely proposed any precise underlying mechanisms. At the same time, research using well-defined multisensory integration paradigms such as the visuotactile ventriloquism aftereffect have rarely explicitly related crossmodal recalibration to body representation. To our knowledge, visuotactile ventriloquism aftereffects have not been viewed as potentially deriving from plasticity in body representations. Here, we directly addressed this potential link by designing a ventriloquism task that allows drawing conclusions about the hypothesized use of one particular body representation, namely one for hand size. Our paradigm extends previous studies with setups that did not allow (and did not aim for) dissociating these different theoretical accounts (Samad & Shams, 2016, 2018).

Our suggestion that recalibration takes place within a bodily map implies that, after experiencing the visuotactile mismatch, tactile stimuli are not just perceived “somewhere else” in space. Instead, recalibration appears to update the representation of how hand landmarks relate to external space and, accordingly, how large the hand is. These updates directly affect both motor actions towards the hand, as demonstrated by the unimodal pointing task, and the spatial perception of stimuli presented anywhere on the hand, as demonstrated by the tactile distance task.

There have been other attempts to change the size representation of the hand using visual stimuli. Examples of this are using large rubber hands in an RHI setup (Filippetti & Crucianelli, 2019; Haggard & Jundi, 2009; Heed et al., 2011), presenting visually resized body parts (Kilteni et al., 2012; Perera et al., 2017; Ratcliffe & Newport, 2017), or resized bodies (van der Hoort et al., 2011) in virtual-reality-like environments. One aspect that all these paradigms have in common is that they aim to resize body representations by showing participants resized bodies or body parts. Notably, the stimuli involved in such paradigms are complex and, accordingly, it is likely that they involve many cognitive processes including low-level processes such as recalibration as well as higher-order conceptual processes related to body features, for example whether the shape or posture of a body part is anatomically possible (Tsakiris, 2010). In contrast, our paradigm uses very basic bimodal pairs of stimuli – dots and taps – that bear no connection to body-related semantics. That a modulation of perceived body size was achieved with this “reduced” paradigm demonstrates that update of body representations does not require any kind of body-shaped stimulus (like a rubber hand) and highlights the role of low-level, bottom-up, multisensory integration and recalibration processes in body perception.

## Conclusions

We report that visual stimuli recalibrate tactile stimuli on the hand such that unimodal tactile stimuli are localized as if the hand had grown. This recalibration also influenced the judgment of tactile distance stimuli on the hand. These body-based visuotactile recalibration effects emerged only when visual and tactile stimuli were spatially congruent and implied that a given bimodal stimulus pair may have originated from a grown hand. This dependence of recalibration on stimulus congruency was not evident in self-reports about the perceptions evoked by the recalibration procedure. We conclude that visuotactile recalibration affects an implicit hand representation that is a common base for both localization and perception of tactile distances.

## Supporting information

Supplementary Analyses

## Acknowledgements

We thank Tine-Marie Wujciak for assistance with piloting and optimizing the paradigm and Franziska Schulz for assistance with data collection. We also thank Luke Miller for recommendations concerning the TDJ procedures.

## References

Alais, D., & Burr, D. (2004). The Ventriloquist Effect Results from Near-Optimal Bimodal Integration. Current Biology, 14(3), 257–262. https://doi.org/10.1016/j.cub.2004.01.029

Barr, D. J., Levy, R., Scheepers, C., & Tily, H. J. (2013). Random effects structure for confirmatory hypothesis testing: Keep it maximal. Journal of Memory and Language, 68(3), 255–278. https://doi.org/10.1016/j.jml.2012.11.001

Bates, D., Mächler, M., Bolker, B., & Walker, S. (2014). Fitting Linear Mixed-Effects Models using lme4. arXiv:1406.5823 [stat]. http://arxiv.org/abs/1406.5823

Bellan, V., Gilpin, H. R., Stanton, T. R., Newport, R., Gallace, A., & Moseley, G. L. (2015). Untangling visual and proprioceptive contributions to hand localisation over time. Experimental Brain Research, 1–13. https://doi.org/10.1007/s00221-015-4242-8

Benjamini, Y., & Hochberg, Y. (1995). Controlling the False Discovery Rate: A Practical and Powerful Approach to Multiple Testing. Journal of the Royal Statistical Society. Series B (Methodological), 57(1), 289–300. https://doi.org/10.2307/2346101

Botvinick, M., & Cohen, J. (1998). Rubber hands „feel” touch that eyes see. Nature, 391(6669), 756. https://doi.org/10.1038/35784

Bruns, P., & Röder, B. (2019). Repeated but not incremental training enhances cross-modal recalibration. Journal of Experimental Psychology: Human Perception and Performance, 45(4), 435–440. https://doi.org/10.1037/xhp0000642

Calzolari, E., Azañón, E., Danvers, M., Vallar, G., & Longo, M. R. (2017). Adaptation aftereffects reveal that tactile distance is a basic somatosensory feature. Proceedings of the National Academy of Sciences, 114(17), 4555–4560. https://doi.org/10.1073/pnas.1614979114

Cardinali, L., Frassinetti, F., Brozzoli, C., Urquizar, C., Roy, A. C., & Farnè, A. (2009). Tool-use induces morphological updating of the body schema. Current Biology, 19(12), R478–R479.

de Vignemont, F., Ehrsson, H. H., & Haggard, P. (2005). Bodily Illusions Modulate Tactile Perception. Current Biology, 15(14), 1286–1290. https://doi.org/10.1016/j.cub.2005.06.067

Ehrsson, H. H. (2007). The Experimental Induction of Out-of-Body Experiences. Science, 317(5841), 1048–1048. https://doi.org/10.1126/science.1142175

Ernst, M. O., & Di Luca, M. (2011). Multisensory perception: From integration to remapping. In J. Trommershäuser, K. Kording, & M. S. Landy (Hrsg.), Sensory cue integration (S. 224–250).

Erro, R., Marotta, A., Tinazzi, M., Frera, E., & Fiorio, M. (2018). Judging the position of the artificial hand induces a “visual” drift towards the real one during the rubber hand illusion. Scientific Reports, 8(1). https://doi.org/10.1038/s41598-018-20551-6

Filippetti, M. L., & Crucianelli, L. (2019). If I were a grown-up: Children’s response to the rubber hand illusion with different hand sizes. Journal of Experimental Child Psychology, 185, 191–205. https://doi.org/10.1016/j.jecp.2019.04.016

Frissen, I., de Gelder, B., & Vroomen, J. (2012). The Aftereffects of Ventriloquism: The Time Course of the Visual Recalibration of Auditory Localization. Seeing and Perceiving, 25(1), 1–14. https://doi.org/10.1163/187847611X620883

Fuchs, X., Riemer, M., Diers, M., Flor, H., & Trojan, J. (2016). Perceptual drifts of real and artificial limbs in the rubber hand illusion. Scientific Reports, 6, 24362. https://doi.org/10.1038/srep24362

Haggard, P., & Jundi, S. (2009). Rubber hand illusions and size-weight illusions: Self-representation modulates representation of external objects. Perception, 38(12), 1796–1803. https://doi.org/10.1068/p6399

Head, H., & Holmes, G. (1911). Sensory Disturbances from Cerebral Lesions. Brain, 34(2–3), 102–254. https://doi.org/10.1093/brain/34.2-3.102

Heed, T., Gründler, M., Rinkleib, J., Rudzik, F. H., Collins, T., Cooke, E., & O’Regan, J. K. (2011). Visual information and rubber hand embodiment differentially affect reach-to-grasp actions. Acta Psychologica, 138(1), 263–271. https://doi.org/10.1016/j.actpsy.2011.07.003

Hirashima, M., & Nozaki, D. (2012). Distinct motor plans form and retrieve distinct motor memories for physically identical movements. Current Biology: CB, 22(5), 432–436. https://doi.org/10.1016/j.cub.2012.01.042

Holmes, N. P., Crozier, G., & Spence, C. (2004). When mirrors lie: „Visual capture” of arm position impairs reaching performance. Cognitive, Affective, & Behavioral Neuroscience, 4(2), 193–200. https://doi.org/10.3758/CABN.4.2.193

Kagerer, F. A., Contreras-Vidal, J. L., & Stelmach, G. E. (1997). Adaptation to gradual as compared with sudden visuo-motor distortions. Experimental Brain Research, 115(3), 557–561. https://doi.org/10.1007/pl00005727

Keizer, A., Smeets, M. A. M., Dijkerman, H. C., van Elburg, A., & Postma, A. (2012). Aberrant somatosensory perception in Anorexia Nervosa. Psychiatry Research, 200(2–3), 530–537. https://doi.org/10.1016/j.psychres.2012.05.001

Kilteni, K., Maselli, A., Kording, K. P., & Slater, M. (2015). Over my fake body: Body ownership illusions for studying the multisensory basis of own-body perception. Frontiers in Human Neuroscience, 9. https://doi.org/10.3389/fnhum.2015.00141

Kilteni, K., Normand, J.-M., Sanchez-Vives, M. V., & Slater, M. (2012). Extending Body Space in Immersive Virtual Reality: A Very Long Arm Illusion. PLoS ONE, 7(7), e40867. https://doi.org/10.1371/journal.pone.0040867

Kuznetsova, A., Brockhoff, P. B., & Christensen, R. H. B. (2014). lmerTest: Tests for random and fixed effects for linear mixed effect models (lmer objects of lme4 package). http://CRAN.R-project.org/package=lmerTest

Lackner, J. R. (1988). Some proprioceptive influences on the perceptual representation of body shape and orientation. Brain, 111(2), 281–297.

Lenggenhager, B., Tadi, T., Metzinger, T., & Blanke, O. (2007). Video Ergo Sum: Manipulating Bodily Self-Consciousness. Science, 317(5841), 1096–1099. https://doi.org/10.1126/science.1143439

Lenth, R., Singmann, H., Love, J., Buerkner, P., & Herve, M. (2019). emmeans: Estimated Marginal Means, aka Least-Squares Means (1.4.3.01). https://CRAN.R-project.org/package=emmeans

Limanowski, J. (2021). Precision control for a flexible body representation. Neuroscience & Biobehavioral Reviews. https://doi.org/10.1016/j.neubiorev.2021.10.023

Linkenhoker, B. A., & Knudsen, E. I. (2002). Incremental training increases the plasticity of the auditory space map in adult barn owls. Nature, 419(6904), Art. 6904. https://doi.org/10.1038/nature01002

Longo, M. R., Azañón, E., & Haggard, P. (2010). More than skin deep: Body representation beyond primary somatosensory cortex. Neuropsychologia, 48(3), 655–668. https://doi.org/10.1016/j.neuropsychologia.2009.08.022

Longo, M. R., & Haggard, P. (2010). An implicit body representation underlying human position sense. Proceedings of the National Academy of Sciences, 107(26), 11727–11732. https://doi.org/10.1073/pnas.1003483107

Longo, M. R., & Haggard, P. (2011). Weber’s illusion and body shape: Anisotropy of tactile size perception on the hand. Journal of Experimental Psychology: Human Perception and Performance, 37(3), 720–726. https://doi.org/10.1037/a0021921

Longo, M. R., & Morcom, R. (2016). No Correlation between Distorted Body Representations Underlying Tactile Distance Perception and Position Sense. Frontiers in Human Neuroscience, 10. https://doi.org/10.3389/fnhum.2016.00593

Mancini, F., Longo, M. R., Iannetti, G. D., & Haggard, P. (2011). A supramodal representation of the body surface. Neuropsychologia, 49(5), 1194–1201. https://doi.org/10.1016/j.neuropsychologia.2010.12.040

Matsumiya, K. (2022). Multiple representations of the body schema for the same body part. Proceedings of the National Academy of Sciences, 119(4), e2112318119. https://doi.org/10.1073/pnas.2112318119

Medina, J., & Coslett, H. B. (2010). From Maps to Form to Space: Touch and the Body Schema. Neuropsychologia, 48(3), 645. https://doi.org/10.1016/j.neuropsychologia.2009.08.017

Medina, J., & Duckett, C. (2017). Domain-general biases in spatial localization: Evidence against a distorted body model hypothesis. Journal of Experimental Psychology: Human Perception and Performance, 43(7), 1430–1443. https://doi.org/10.1037/xhp0000397

Miller, L. E., Cawley-Bennett, A., Longo, M. R., & Saygin, A. P. (2017). The recalibration of tactile perception during tool use is body-part specific. Experimental Brain Research, 235(10), 2917–2926. https://doi.org/10.1007/s00221-017-5028-y

Miller, L. E., Longo, M. R., & Saygin, A. P. (2014). Tool morphology constrains the effects of tool use on body representations. Journal of Experimental Psychology: Human Perception and Performance, 40(6), 2143–2153. https://doi.org/10.1037/a0037777

Miller, L. E., Longo, M. R., & Saygin, A. P. (2016). Mental body representations retain homuncular shape distortions: Evidence from Weber’s illusion. Consciousness and Cognition, 40, 17–25. https://doi.org/10.1016/j.concog.2015.12.008

Oldfield, R. C. (1971). The assessment and analysis of handedness: The Edinburgh inventory. Neuropsychologia, 9(1), 97–113.

Perera, A. T., Newport, R., & McKenzie, K. J. (2017). Changing hands: Persistent alterations to body image following brief exposure to multisensory distortions. Experimental Brain Research, 1–13. https://doi.org/10.1007/s00221-017-4935-2

Pinheiro, J. C., & Bates, D. M. (2000). Mixed-effects models in S and S-PLUS. Springer.

Ratcliffe, N., & Newport, R. (2017). The Effect of Visual, Spatial and Temporal Manipulations on Embodiment and Action. Frontiers in Human Neuroscience, 11. https://doi.org/10.3389/fnhum.2017.00227

Recanzone, G. H. (1998). Rapidly induced auditory plasticity: The ventriloquism aftereffect. Proceedings of the National Academy of Sciences of the United States of America, 95(3), 869–875. https://doi.org/10.1073/pnas.95.3.869

Riemer, M., Trojan, J., Beauchamp, M., & Fuchs, X. (2019). The rubber hand universe: On the impact of methodological differences in the rubber hand illusion. Neuroscience & Biobehavioral Reviews, 104, 268–280. https://doi.org/10.1016/j.neubiorev.2019.07.008

Samad, M., & Shams, L. (2016). Visual-somatotopic interactions in spatial perception. Neuroreport, 27(3), 180–185. https://doi.org/10.1097/WNR.0000000000000521

Samad, M., & Shams, L. (2018). Recalibrating the body: Visuotactile ventriloquism aftereffect. PeerJ, 6, e4504. https://doi.org/10.7717/peerj.4504

Schönbrodt, F. D., & Perugini, M. (2013). At what sample size do correlations stabilize? Journal of Research in Personality, 47(5), 609–612. https://doi.org/10.1016/j.jrp.2013.05.009

Tamè, L., Azañón, E., & Longo, M. R. (2019). A Conceptual Model of Tactile Processing across Body Features of Size, Shape, Side, and Spatial Location. Frontiers in Psychology, 10. https://doi.org/10.3389/fpsyg.2019.00291

Tosi, G., Maravita, A., & Romano, D. (2022). I am the metre: The representation of one’s body size affects the perception of tactile distances on the body. Quarterly Journal of Experimental Psychology, 75(4), 583–597. https://doi.org/10.1177/17470218211044488

Tsakiris, M. (2010). My body in the brain: A neurocognitive model of body-ownership. Neuropsychologia, 48(3), 703–712. https://doi.org/10.1016/j.neuropsychologia.2009.09.034

van der Hoort, B., Guterstam, A., & Ehrsson, H. H. (2011). Being Barbie: The Size of One’s Own Body Determines the Perceived Size of the World. PLoS ONE, 6(5), e20195. https://doi.org/10.1371/journal.pone.0020195

Wann, J. P., & Ibrahim, S. F. (1992). Does limb proprioception drift? Experimental Brain Research, 91(1), 162–166.

Wickham, H. (2009). ggplot2: Elegant graphics for data analysis. Springer Science & Business Media.

