## Supplementary Analyses for "Modulation of implicitly perceived hand size by visuotactile recalibration"

### Modulation of implicitly perceived hand size by visuotactile recalibration—Supplementary material

#### 1 Intensity Ratings

We assessed stimulus intensity ratings for the tactile stimulators at all 8 stimulus locations to ensure that participants indeed perceived all stimuli. Participants judged stimulus intensity, on average, between 3.6 (D5 knuckle; maximum rating: 8) and 6.1 (D1 tip). This shows that the stimuli were perceived at a moderate intensity.

Table S1. Average (mean) and standard deviations for the intensity ratings at the 8 hand landmarks.

| Landmark | Mean | SD |
|---|---|---|
| B | 4.4 | 2.2 |
| D1 | 6.1 | 2.2 |
| D2 | 5.3 | 2.1 |
| D3 | 4.9 | 2.0 |
| D4 | 4.0 | 1.9 |
| D5 | 4.2 | 2.0 |
| D2K | 4.4 | 1.7 |
| D5K | 3.6 | 1.7 |

#### 2 Coefficients of linear and quadratic components of the localization map size change over the recalibration iterations

Here we present the coefficients of the model presented in the "Size increase in the localization maps" of main text.

##### 2.1 Linear component

| congruency | *b* linear trend | SE | DF | lower CL | upper CL | t ratio | p | sign. |
|---|---|---|---|---|---|---|---|---|
| congruent | 1.198 | 0.27 | 29.77 | 0.65 | 1.75 | 4.44 | <0.001 | *** |
| incongruent | 0.586 | 0.27 | 29.77 | 0.03 | 1.14 | 2.17 | 0.038 | * |

##### *2.2 Quadratic component*

| congruency | *b* quadratic trend | SE | DF | lower CL | upper CL | t ratio | p | sign. |
|---|---|---|---|---|---|---|---|---|
| congruent | -0.806 | 0.22 | 32.99 | -1.25 | -0.36 | -3.66 | 0.002 | ** |
| incongruent | -0.323 | 0.22 | 32.98 | -0.77 | 0.12 | -1.47 | 0.152 | |

#### *3 Length vs. width anisotropies in tactile distance judgments*

Tactile distances were judged significantly larger when the stimuli were applied in the mediolateral as compared to the proximodistal direction. This was evident from a main effect of *direction* in the LMM ($F(1, 24) = 65.07$, $p < 0.001$). As shown in Figure 6B in the main text, participants underestimated distance in the proximodistal and overestimated it in the mediolateral direction. The difference between directions depended on the stimulus distance and was larger for the 3.3 and 4.2 cm distances as compared to the 2.4 cm distance (*direction* × *distance* interaction: $F(2,7907) = 102.26$, $p < 0.001$; see Table 5). However, post-hoc comparisons revealed that the anisotropies (differences between mediolateral and proximodistal directions) were significant for each of the three distances (see Table S2).

Table S2: Post-hoc comparisons between tactile distance judgements for the three distance stimuli in proximodistal and mediolateral direction.

| distance | contrast | estimate | SE | DF | t ratio | lower CL | upper CL | p | sign. |
|---|---|---|---|---|---|---|---|---|---|
| 2.4 | proximodistal - mediolateral | -0.62 | 0.14 | 27.08 | -4.29 | -0.91 | -0.32 | <0.001 | *** |
| 3.3 | proximodistal - mediolateral | -1.32 | 0.14 | 27.08 | -9.14 | -1.61 | -1.02 | <0.001 | *** |
| 4.2 | proximodistal - mediolateral | -1.44 | 0.14 | 27.08 | -10.01 | -1.74 | -1.15 | <0.001 | *** |
